# Predicting Coexistence in Species with Continuous Ontogenetic Niche Shifts and Competitive Asymmetry

**DOI:** 10.1101/119446

**Authors:** Ronald D. Bassar, Joseph Travis, Tim Coulson

**Author notes:** Corresponding author –. Statement of authorship: RDB, JT, and TC conceived the ideas presented. RDB developed and analyzed the models with input from JT and TC. RDB wrote the first draft of the manuscript and all authors contributed substantially to the final version.

## Abstract

A longstanding problem in ecology is whether structured life cycles impede or facilitate coexistence between species. Theory based on populations with two discrete stages in the life-cycle indicates that coexistence requires at least one species to shift its niche between stages and that each species is a better competitor in one of the niches. However, in many cases, niche shifts are associated with changes in an underlying continuous trait like organism size and we have few predictions for how the conditions for coexistence are affected by this type of ontogenetic dynamics. Here we develop a framework for analyzing species coexistence based on Integral Projection Models (IPMs) that incorporates continuous ontogenetic changes in both the resource niche and competitive ability. We parameterize the model using experimental data from Trinidadian guppies and make predictions about how niche shifts and competitive symmetries allow or prevent species coexistence. Overall, our results show that the effects of competition on fitness depend upon trait-mediated niche-separation, trait-mediated competitive asymmetry in the part of the niche that is shared across body sizes, and the sensitivity of fitness to body size. When all three conditions are considered, we find multiple ecological and evolutionary routes to coexistence. When both species can shift their niche with increasing body size and competition for resources among the species and sizes is symmetric, then the species that shifts its niche to a greater degree with ontogeny will competitively exclude the other species. When competitive ability increases with increasing body size, then the two species can coexist when the better competitor shifts its niche with body size to a lesser degree than the weaker competitior. This region of coexistence shrinks as the better competitor increasingly shifts its niche with increasing size. When both species shift their niches with size, but each is a better competitor on resources used by smaller or larger individuals, then the model predicts an alternative stable state over some range of niche shifts. We discuss how our results provide new insights into species coexistence and the evolutionary consequences of size-structured interspecific competition.

## INTRODUCTION

For many organisms, resource use patterns change throughout development (Werner and Gilliam 1984). Such ontogenetic niche shifts range from abrupt changes that delineate discrete life stages as seen in amphibians or many insects to continuous changes in niche use as a function of a trait linked to development (e.g. body size in many fishes, mammals, and reptiles). Some organisms exhibit both discrete and continuous niche shifts. For example, dragonflies occupy discrete niches with aquatic naiads and terrestrial adults but the food habits of the naiads change dramatically as size and instar number increases. Such ontogenetic changes lead to the structuring of populations into both discrete and continuous stages where individuals in one stage may compete with individuals in the same stage and in different stages or among different sizes in contrasting ways.

One persistent question associated with such structured life cycles is whether they promote or impede species coexistence (Miller and Rudolf 2011, Nakazawa 2015). In simple, two-species models with two discrete life stages (e.g. larvae and adults), coexistence of two species depends on whether one, or both, species shift their niche completely with stage, and which species is a better competitor in each of the two stages (Haefner and Edson 1984, Loreau and Ebenhoh 1994, Moll et al. 2008, Ackleh and Chiquet 2011, de Roos and Persson 2013). In these cases, the niche shift is discrete and there is little or no niche overlap between the stages. When one species shifts completely and the other does not, coexistence is possible if the species that does not shift its niche is a slightly better competitor than the one that does shift its niche (e.g. Loreau and Ebenhoh 1994, de Roos and Persson 2013). For coexistence to occur when both species shift their niches completely with stage, each species must be a better competitor in one of the stages, a worse competitor in the other stage, and be limited in the stage in which it is the better competitor (e.g. Loreau and Ebenhoh 1994, McCann 1998, de Roos and Persson 2013). In cases where this is not true, there will be either deterministic competitive exclusion by one species or a priority effect where the species that initially occupies the habitat prevails.

While these predictions apply to many life histories, for a considerable number of species, life stages cannot be categorized into discrete stages. For these species, the niche shift is more accurately described by continuous changes in a trait that is related to niche use and competitive asymmetry (Werner and Gilliam 1984). Competitive asymmetry means that some individuals are better able to gather resources or are more efficient at utilizing the resources for organismal functions (somatic growth or reproduction). For a wide variety of organisms, the trait that determines the niche and competitive ability is body size. It is unclear how coexistence theory built on niche shifts between discrete life stage changes translates into predictions of coexistence for organisms with continuous life stages (Nakazawa 2015).

Many of the insights that we have about coexistence in size-structured populations are derived from elegant models of the dynamics of two or more consumers competing for two or more resources (Haefner and Edson 1984, Loreau and Ebenhoh 1994, McCann 1998, Moll et al. 2008, Ackleh and Chiquet 2011, de Roos and Persson 2013). In this approach, the dynamics of the resources are explicitly modelled, which allows predictions about how coexistence is influenced by those resource dynamics or how coexistence conditions change along gradients of productivity (e.g. de Roos and Persson 2013). These models produce predictions when juveniles and adults consume completely different resources. When juveniles and adults overlap in resource use to varying degrees, the predictions are often contingent upon the relative values of several parameters that can be very difficult to estimate empirically. This makes it difficult to test those predictions outside of simple laboratory systems where resource dynamics can be controlled experimentally.

An alternative approach to this challenge is to use simplified models that capture the essence of the conceptual issue, but that can be parameterized using observational or experimental data frequently collected by ecologists. The degree to which these approaches agree can then be seen as a path for validating the simpler approach. We take that approach here, developing a framework for analyzing coexistence in discrete time models with continuous traits that determine ontogenetic changes in niche use and competitive asymmetry. We develop a general, single-sex, two-species Integral Projection Model (IPM) where each species can shift its niche to varying degrees with development. Atop this model we overlay asymmetries in competition that depend on body size. We then use invasion analyses to ask how asymmetries influence predictions about species coexistence. Finally, we use perturbation analysis of the model to describe how different life stages contribute to the transition between competitive exclusion and species coexistence. We then discuss how this framework produces shared or unique predictions about coexistence compared to previous models of coexistence in structured populations.

## MATERIALS AND METHODS

### MODELLING FRAMEWORK

We follow the structured modelling approach developed in our previous work on competitive asymmetries based on Integral Projection Models (IPMs) (Bassar et al. 2016). A simple single species, single sex (female only) IPM is written:

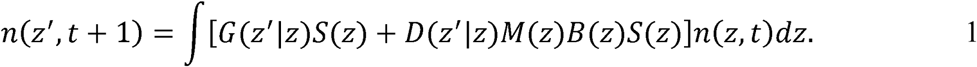

The function *n*(*z*,*t*) describes the number of individuals in the population at time *t* across all *z* trait values such that ∫ *n*(*z*,*t*)*dz* is the total number of individuals in the population. Here we assume *z* is body length, but the model can be used to describe any continuous trait that changes with ontogeny. *S*(*z*) and *B*(*z*) are functions respectively describing the probability of an individual with body size *z* surviving the interval and reproducing at the end of the interval. *M*(*z*) is a function describing the expected litter size of an individual with body size *z*. *G*(*z*′|*z*) describes the probability of a surviving individual with body size *z* at time *t* growing to body size *z′* at time *t* + *1. D*(*z′*|*z*) describes the probability that a parent with body size *z* at time *t* produces an offspring with trait value *z*′ at time *t* + 1. The structure of the model assumes that individuals give birth to offspring immediately preceding the next census.

The relationship between size and each of these vital rates can be estimated using generalized regression methods (Easterling et al. 2000, Coulson et al. 2010, Coulson 2012, Rees et al. 2014, Bassar et al. 2016). For example, the probability of surviving the interval can be estimated using a generalized linear model with a binomial error structure and logit link function:

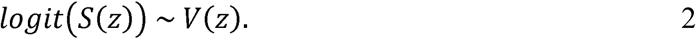

Equations for each of the vital rates are given in Table 1.

**Table 1.**
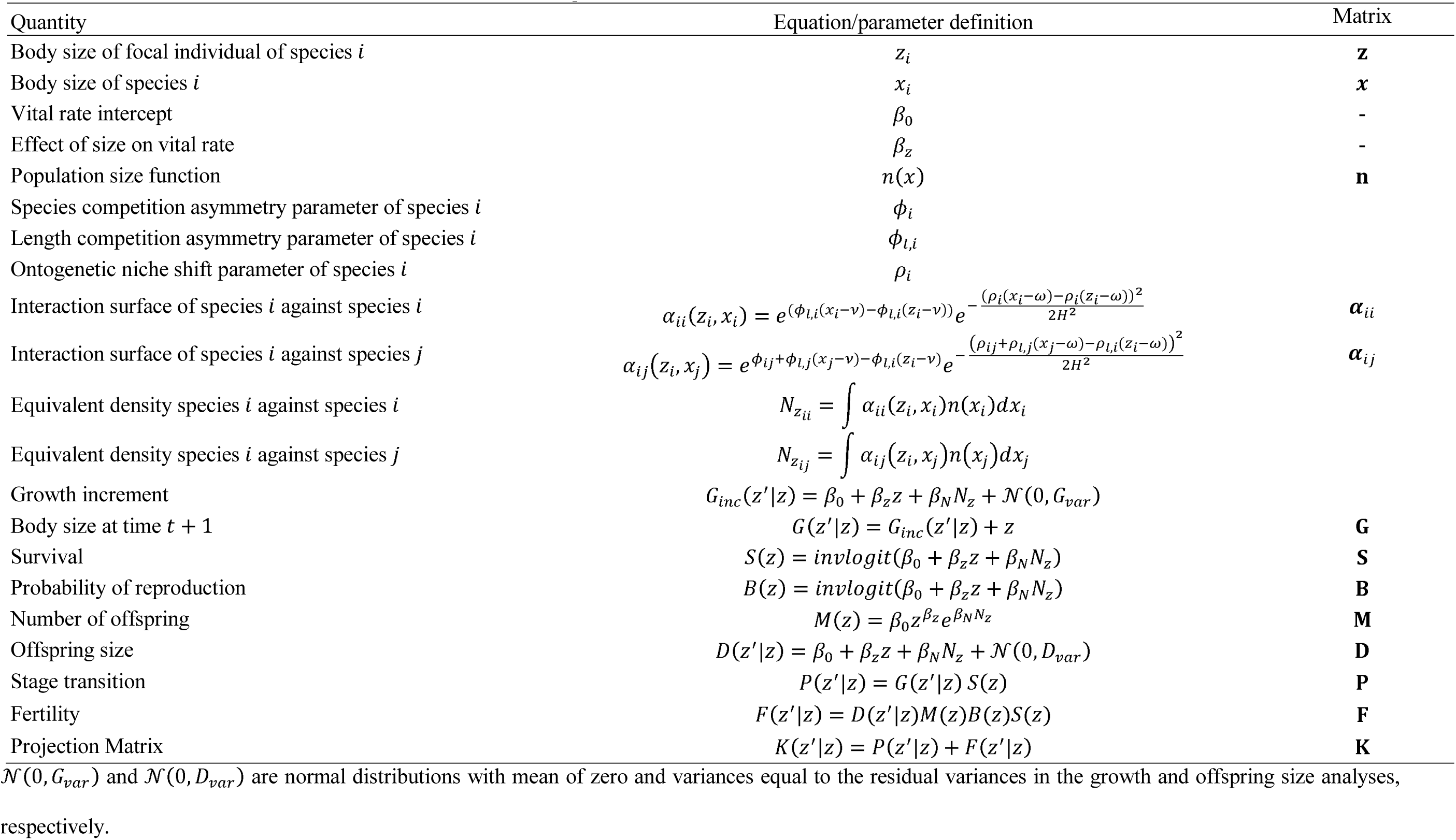
Parameters definitions, vital rate functions and matrix equivalents.

Bassar et al. (2016) showed that one way to incorporate density- and frequency-dependence is to make the linear function describing any vital rate, *V*, a function of body size and the number and distribution of body sizes (or other traits) in the population as:

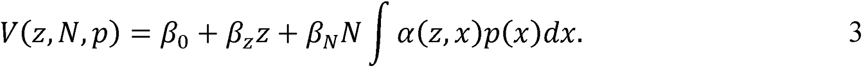

Here *z* represents the trait value of a focal individual and *x* represents the trait value of a competitor within the population. *N* is the total population density and *p*(*x*) = *N*^−1^∫*n*(*x*)*dx* is the frequency of individuals with trait *x*. The terms, *β*_0_ + *β_z_z*, describe the expected value of the vital rate when *N ∫p*(*x*)*dx* = 0 (i.e. the density-independent terms) and how it changes as a function of body size, *z*. The parameter *β_N_* describes how the vital rate is affected by a change in the density of the population, independent of the trait values in the population. When density decreases the vital rate, as in competitive interactions, *β_N_* is negative. The function *α*(*z*,*x*) is an interaction surface. It describes the strength of competitive interactions, measured as the number of individuals with trait value *x* that are equivalent to an individual with trait value *z* (Bassar et al. 2016). Bassar et al. (2016) describe how asymmetric competition between sizes in single species models alters the vital rates, population dynamics, and evolutionary quantities such as generation time and provide alternative formulations of the vital rate equations that are not additive.

Extending the model to more than one species involves adding an additional term to the vital rate equations that describe the impact of the number of size *x* individuals of another species. If *V_i_* is the vital rate equation for species *i, z_i_* is the body size of a focal individual of species *i*, and *x_j_* represents the body sizes of species *j*, then our equation for the vital rate of species *i* is:

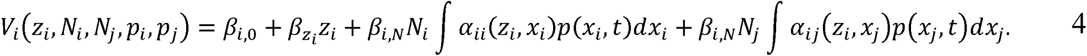

The interaction surface *α_ii_*(*z_i_*,*x_i_*) describes the within species equivalence of an individual of with body size *x* on an individual with body size *z*. The interaction surface *α_ij_*(*z_j_*,*x_j_*) describes the between species equivalence of an individual of species *j* with body size *x* on an individual of species *i* with body size *z*.

The interaction surface can have many forms, depending on how resources are partitioned. Bassar et al. (2016) describe several possible forms when competitive asymmetries among individuals depend on body size, but where all individuals are assumed to compete for the same resources. Interaction surfaces _that_ incorporate ontogenetic niche shifts, competitive asymmetries between species, and competitive asymmetries between body sizes have a different form. Following Macarthur and Levins (1967), we assume that the distribution of resource utilization of species *i* can be described by the equation 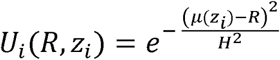, where *μ*(*z_i_*) is the mean position of the distribution on a linearized resource axis, *R*, and *H^2^* is the niche width, which we assume to be equal to 1 for all sizes and species. We assume the mean niche position is a linear function of body size such that *μ*(*z_i_*) = *ρ_i_* + *ρ_l,i_*(*z_i_* − *ω*). The parameter *ρ_i_* defines the mean niche position of body size *ω* in species *i* and *ρ_l,i_* describes how the mean niche position changes with body size. We then assume that the area under the niche function can be interpreted as the total resource utilization and can be scaled by a linear function of body size on the natural log scale *e*^*ϕ_i_+ϕ_l,i_*(*z_i–v_*)^, where the parameter *ϕ_i_* defines the resource use of body size *v* in species *i* and *ϕ_l,i_* describes how resource acquisition changes with body size. Combining these functions yields: 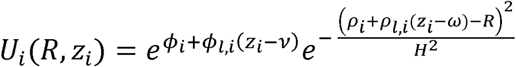. In keeping with the standard interpretation of interaction coefficients as relative effects, the interaction surface, or the amount of niche overlap scaled by competitive asymmetry between species is then calculated as:

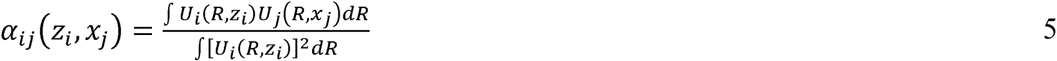

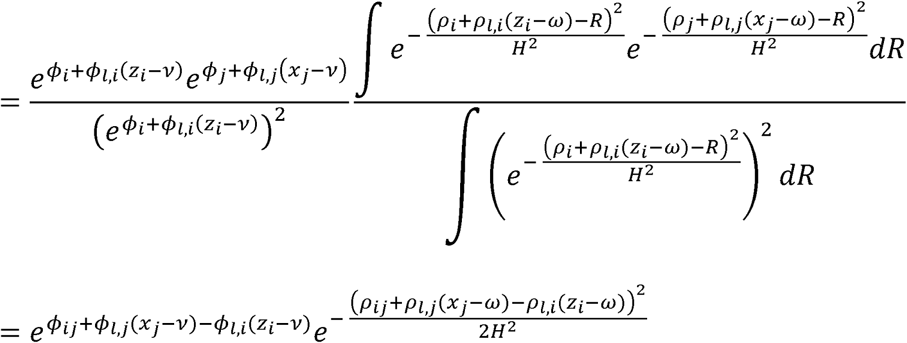
where *ρ_ij_* is the difference in the mean niche positions between species at size *ω* and *ϕ_ij_* measures the difference in resource use between the species at size *v*.

Each of the parameters of the interaction surface alters the shape of the interaction surface in different ways (Figure 1). When species *i* does not shift its niche with size (*ρ_i_* = 0), species *j* shifts to a large degree (*ρ_j_* = 0.15) and all individuals of both species are equal competitors (symmetric interaction), then the intraspecific interaction surface for species *i* is flat (row 1, column 1 of Figure 1). In contrast, the interspecific interaction surface decreases with increasing competitor size (row 1, column 2 of Figure 1). For species *j*, the interspecific interaction surface decreases with increasing size of individuals of species 2 (row 1, column 3 of Figure 1). The intraspecific interaction surface of species 2 leads to a situation where individuals of species *j* compete most strongly with individuals of the same size (row 1, column 4 of Figure 1). Competition then decreases as the difference in size between the focal individual and the competitor increases (row 1, column 4 of Figure 1). Increasing the competitive advantage of larger individuals of species *i*, then distorts the shape of all interaction surfaces involving species such that the competitive effect of larger individuals of species *i* have greater effects on competition (row 2 of Figure 1).

**Figure 1.**
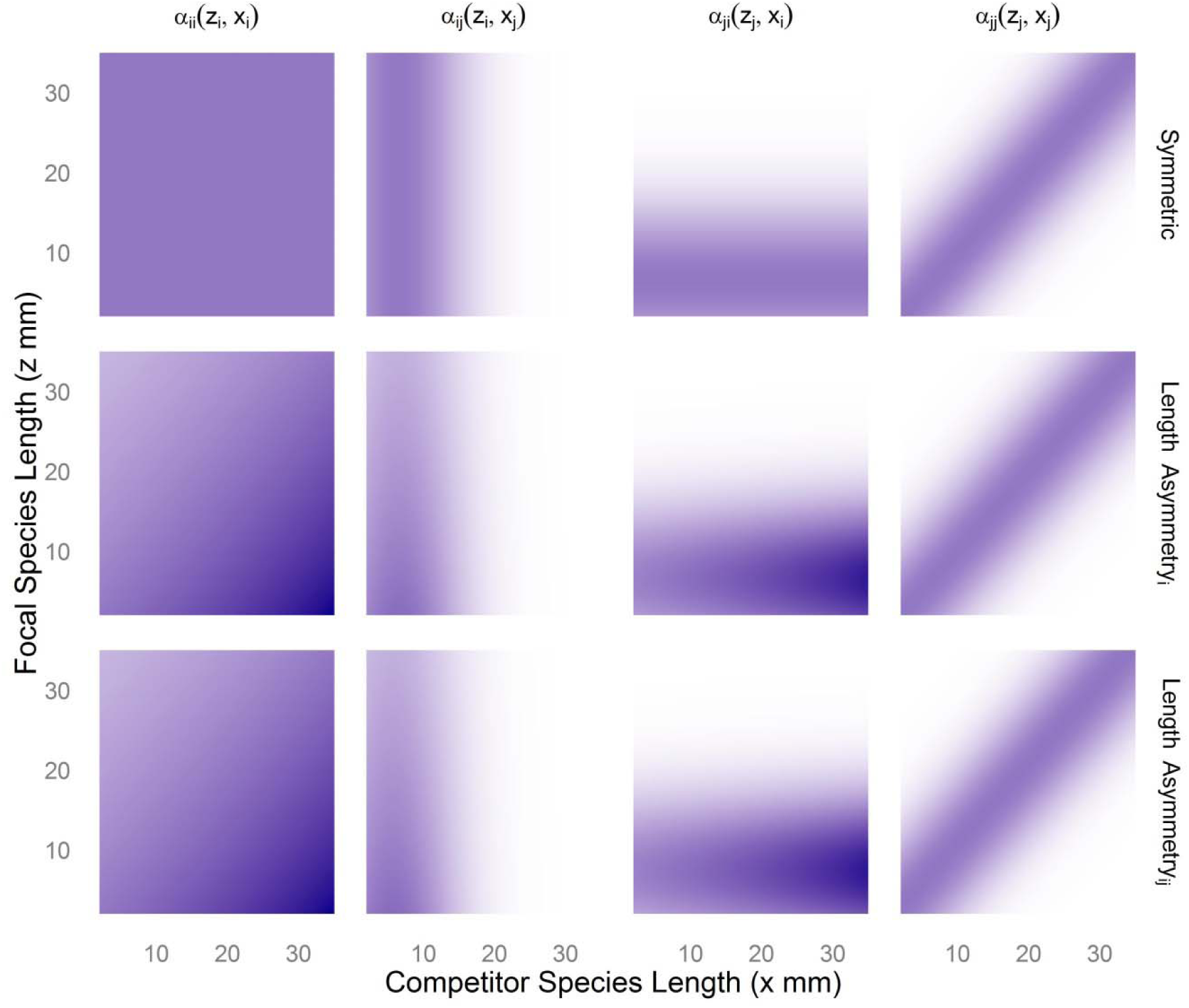
Interaction surfaces for two species model when species *i* does not shift its niche with size *ρ_i_* = 0 and species *j* does (*ρ_j_* = 0.15). Columns give the interaction surfaces for the within and between species interactions for species and the between and within species interactions for species *j*, respectively. Rows represent the three ecological scenarios presented: all individuals of both species are equally competitive (symmetric), competitive ability increases with body length (length asymmetry), and competitive ability increases with length in species and decreases with length in species *j* (length asymmetry). White space represents regions where the interaction surface is near or equal to 0. Violet areas are where the interaction surface is equal to 1, as in the upper left panel. Dark blue areas are regions where the interactions surfaces are greater than 1.

## MODEL PARAMETERIZATION

To facilitate the analysis of the model under realistic conditions, we parameterized our model using demographic data from Trinidadian guppies (*Poecilia reticulata*). Guppies are small stream fish that inhabit freshwater streams on the Caribbean island of Trinidad. In our analysis, we assume both species in the model have an identical guppy life history—the *β* parameters in equation 4 are the same between the species. The model is parameterized from the results of experiments in mesocosms and in natural streams aimed at parameterizing density-dependent IPMs. Briefly, each of these experiments used guppies from streams where they and Hart’s killifish (*Rivulus hartii*) are the only fish species (i.e. the so-called “low predation” guppy life history). In the mesocosm experiments, guppies were added at two densities. In the real stream experiments, natural populations were either decreased by one-half or kept at ambient densities. The duration of each experiment was 28 days, which is slightly longer than one reproductive cycle of guppies. At the end of the mesocosm experiments, guppies were removed from the mesocosms and measured for somatic growth (mm standard length, hereafter SL), reproductive status (pregnant or not pregnant), fecundity, and offspring size (mm SL). Similar measurements were taken from the fish in the experiments in natural streams, but we only use data on survival here. More details of the experiments are described in detail in Bassar et al. (2013). The parameters were estimated using general linear mixed or generalized linear mixed models assuming guppies do not shift their niche with size (*ρ* = 0) and that competition among sizes is symmetrical (*ϕ_i_* = 0) (see Table 2 for estimated parameters). As in Bassar et al. (2016), we did not attempt to fit values for *ρ* or *ϕ* because this requires more detailed experiments than are currently available. The interval over which the IPM projects population dynamics is 28 days. The size interval over which the projection is calculated is 2 mm to 35 mm standard length (SL). Although guppies are never smaller than 5 mm at birth, decreasing the lower limit prevents any loss of individuals from the modelled population (Williams et al. 2012). Likewise, wild guppies from these populations rarely obtain 30 mm in size and the extreme upper limit prevents the unintentional loss of adults from the population (Williams et al. 2012).

**Table 2.** Parameters and standard errors from mesocosm and field studies of Trinidadian guppies.

| Parameter | Mean Growth |  | Survival |  | Fecundity |  | Prob. of Repro |  | Offspring Size |  |
| --- | --- | --- | --- | --- | --- | --- | --- | --- | --- | --- |
|  | Est | SE | Est | SE | Est | SE | Est | SE | Est | SE |
| $\beta_0$ | 3.55 | 0.428 | 0.94 | 0.239 | 2.06 | 0.215 | 2.86 | 1.238 | 6.69 | 0.691 |
| $\beta_z$ | -0.31 | 0.015 | 0.06 | 0.027 | 2.32 | 0.504 | 0.60 | 0.106 | - | - |
| $\beta_N$ | -0.12 | 0.031 | -0.03 | 0.071 | -0.05 | 0.015 | -0.13 | 0.079 | 0.02 | 0.017 |
| Stage of Development | - | - | - | - | - | - | - | - | -0.01 | 0.007 |
| Mesocosm | 0.16 | 0.395 | - | - | - | - | 0.17 | 0.417 | - | - |
| Drainage | - | - | - | - | - | - | 0.04 | 0.193 | 0.80 | 0.896 |
| Residual Variance | 0.45 | 0.669 | - | - | - | - | - | - | 0.17 | 0.412 |

## MODEL ANALYSIS

We begin by describing three general ecological scenarios where one or both species display different competitive asymmetries and ontogenetic niche shifts. We then describe how we derived predictions about whether the scenarios predict coexistence or competitive exclusion using invasion analyses. We next describe how we analyzed the boundary conditions between these two predictions. Finally, we use a sensitivity analysis to ask how small perturbations of the parameters describing the interaction surface cause size-specific effects that lead to coexistence or competitive exclusion.

### Ecological scenarios and predictions of coexistence and competitive exclusion

In scenario 1, we assumed that individuals of both species shifted their niche to varying degrees (*ρ*_*l*,1_ ≥ 0, *ρ*_*l*,2_ ≥ 0) and that there were no size based competitive asymmetries (*ϕ*_*l*,1_ = 0,*ϕ*_*l*,2_ = 0). For this and all other scenarios, we assumed that there were no general differences in the niches between species (*ρ_ij_* = 0) and that each species had equal competitive ability (*ϕ_ij_* = 0). We initiated these scenarios using niche shift (*ρ_l_*) values for both species that range from 0 (no ontogenetic shift) to 0.17, a value at which the largest individuals do not compete with the smallest individuals. We chose to use *v* = 6.5 and *ω* = 6.5 so that in the case of no ontogenetic niche shifts of species *i*, all individuals of species *i* compete most strongly with newborns of species *j* (which are ∼ 6.5 mm) and vice versa. Throughout this range of *ρ_i_* values, the resident population always converged upon a single, stable equilibrium distribution (determined through iterating the resident population through time until *λ* = 1).

In scenario 2 we modelled an ecological situation where the competitive ability of individuals of species 1 increased with increasing body size and, as in scenario 1, each species could shift its niche to varying degrees. The increases in competitive ability of larger individuals of species 1 were modelled by changing *ϕ_l_*_,1_ from 0 to 0.02. When *ϕ_l_*_,1_ = 0.02, a 20mm individual of species 1 has a competitive advantage over a 10mm individual of species 1 of about 22% and all individuals of species 2 of about 49%. As in scenario 1, we held *υ* = 6.5 and *ω* = 6.5. The consequence of this decision was that mean size newborns of both species occupied the same niche and were equally competitive. Competitive ability of species 2 remained the same throughout life, but competitive ability of species 1 increased with increasing size.

In scenario 3, we modelled an ecological situation where larger individuals of species 1 are better competitors and smaller individuals of species 2 are better competitors and, as in scenarios 1 and 2, each species could shift its niche to varying degrees. We changed *ϕ*_*l*, 1_ from 0 to 0.06 and *ϕ*_*l*, 2_ from 0 to -0.06. We set *υ* = 15 and *ω* = 6.5. The consequence of these parameter values are that below 15mm, individuals of species 1 are worse competitors than individuals of the same size of species 2, but above 15mm individuals of species 1 are better competitors than individuals of species 2 of the same size. Overall, this last scenario represents the case where each species is a better competitor at different life stages.

For each of the ecological scenarios, the parameters related to competitive asymmetry (*ϕ*_21_, *ϕ*_*l*_) or ontogenetic niche shifts (*ρ*) were perturbed in all of the demographic rates. Thus, the scenarios do not represent scenarios where changes in competitive asymmetry influences, for example, somatic growth rate but not the other demographic rates.

### Coexistence as an invasion analysis problem

Coexistence means that both species can occupy a habitat despite, or perhaps because of, interspecific interactions. Historical approaches to understanding the conditions for species coexistence involved asking whether there exists a stable equilibrium where all or a subset of the species had population sizes greater than zero. Although this criterion is useful when the dynamics of the system represent fixed-point attractors (stable equilibrium), it is less tractable when the populations are structured (e.g. by age or size) or when the attractors are not fixed points (e.g. periodic, cycles, chaotic). Modern concepts use an approach based on the examining the stability of the trivial equilibrium (where: 0 = *N_i_* = *∫ n*(*X_i_, t*)*dx_i_*) of one species (the invader) while the other species (the resident) resides on its single species attractor. A similar approach is used in calculating evolutionary stable strategies (ESS) in evolutionary biology. Coexistence is predicted to occur when the trivial equilibrium of both species is unstable when evaluated in the context of the other species on the attractor of the single species (or clone in ESS analyses). When the trivial equilibrium of the invader is unstable, then the population should move towards a positive attractor. The interpretation of an unstable equilibrium under these conditions is that each species could grow when rare (Roth et al. 2016). When the trivial equilibrium of the invader is stable, then the population should return towards *N_i_* = 0. Such an approach is valid for a wide range of population and community dynamics including those that are chaotic (Roth et al. 2016).

The stability of the trivial equilibrium is evaluated using the typical tools of a stability analysis, which involve making small perturbations to the invader population size and evaluating its growth rate. If species *i* is the invader and species *j* the resident, then the first-order matrix approximation of this perturbation for IPMs at the trivial equilibrium of the invader is:

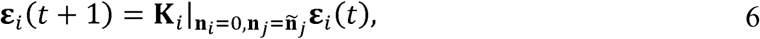
where ***ε**_i_*(*t*) is a vector containing values that represent the density of invading individuals of species *i* and 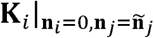 is the projection matrix approximation of the continuous projection kernel. It is a square projection matrix with row elements denoted *r* and column elements denoted *s*. It is evaluated at the trivial equilibrium of species *i* and the positive attractor of species *j* (see Caswell 2001 and the Online Supplement for details). In the simplest case, the attractor is a stable, fixed equilibrium population distribution, **ñ**_*j*_, but this need not be the case (Caswell 2001). Here we discuss the case where the attractor of the resident species is a stable, fixed equilibrium distribution. Stability of the trivial equilibrium of species *i* is then given by the dominant eigenvalue (*λ_inv_ij__*) of the matrix 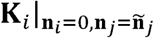. The trivial equilibrium is stable if *λ_inv_ij__* < 1 and unstable if *λ_inv_ij__* > 1. More generally, coexistence then requires mutual invasibility between the species. In other words, species *i* can invade species *j* (*λ_inv_ij__* > 1) and species *j* can invade species *i* (*λ_inv_ij__* > 1).

#### Analysis of the boundaries between coexistence and competitive exclusion

We then present an analysis of the boundaries between competitive exclusion and coexistence. This analysis involves first calculating the values of the parameters (*ρ_l_* and *ϕ_l_*) where *λ*_*inv*_21__ = 1 or *λ*_*inv*_12__ = 1. We calculated these values of the parameters using numerical iteration under a situation where species 1 does not shift its niche with size and species 2 shifts it niche to varying degrees in scenarios 1, 2 and 3. In scenario 3, we additionally calculated these boundaries assuming that both species shift their niches equally with size. This was done so that we could capture the boundary of the priority effects predicted in this scenario. At these points, the invasion growth rates are equal to the growth rate at the single species equilibrium (*λ_inv_ij__* = *λ_ij_* = 1), meaning that the effect of intraspecific competition on the population growth rate at the single species equilibrium is equal to the effect of interspecific competition on the invasion growth rate. The effect of competition on the growth depends on the strength of competition across sizes (which itself is a function of size-dependent niche shifts and size-dependent competitive ability) and, critically, the contribution of the different sizes to the growth rate of the population. We present the strength of intra and interspecific competition experienced at the boundary between coexistence and exclusion as:

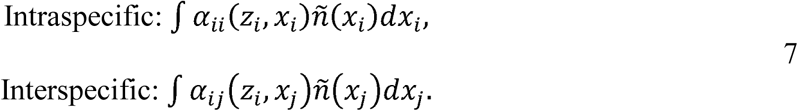

The interpretation of these functions is the equivalent density of individuals of species *i* or *j* experienced by an individual of species *i* with size *z*. It is important to note that neither these functions nor their integrals with respect to *z* represent effects on either growth rate and will not necessarily be equal to each other at the boundary. What *will* be equal at the boundary are the *effects* of each type of competition on *λ*. We present these functions to show the strength of competition acting at the range of sizes at the boundary.

#### Sensitivity analyses at the boundary between coexistence and competitive exclusion

We then used a sensitivity analysis of *λ_inv_* with respect to the model parameters to assess how increases in competitive asymmetry between sizes promote coexistence or competitive exclusion through the different sizes. In the Online Supplement, we show that the matrix approximation of this sensitivity is:

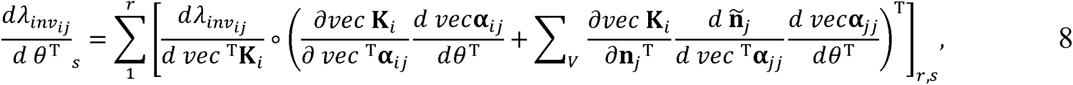
where *θ* is a scalar that represents any of the parameters in the model. Importantly, **K_*i*_** is constructed at the single species equilibrium of the resident (species *j*) and the trivial equilibrium of the invader (species *i*). The operator °, means to take the element by element product and *vec* means to vector transform the matrix so that *vec* **K**_*i*_ = [*k*_1,1_, …, *k*_*r*,1_, *k*_1,2_, …, *k*_*r*,2_, …, *k*_1,*s*_, …, *k*_*r*,*s*_]^T^, or so that columns, *s*, are stacked one on top of the other. The derivative, 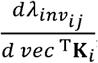, is the vector transformed sensitivity of the invasion growth rate of species *i* to changes in the elements of the matrix approximation of the IPM projection kernel. The product of this derivative and the first term in parentheses describes the direct effect the parameter has on the invasion growth rate of the invader. The product of 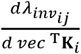 and the second term in parentheses describes how invasion of species *i* changes through the effect that *ψ* has on the equilibrium distribution of the resident. We present these size dependent sensitivities to show how small changes in the parameters influences invasion across the range of body sizes.

## RESULTS

### ECOLOGICAL SCENARIO 1: ONTOGENETIC NICHE SHIFTS WITH COMPETITIVE SYMMETRY

Coexistence between two species is not possible when interspecific competition is symmetric (*ϕ*_*l*_ = 0) and one species shifts its niche with size more than the other (Figure 2A). Instead, the species that shifts its niche with ontogeny to a greater degree will exclude the other species (Figure 2A). When both species shift niches with size to an equal degree (diagonal line in Figure 2A), the invasion growth rate (*λ_inv_*) of both species is equal to 1 (represented by the black dot in Figure 2B) and coexistence is neutral. When this condition is met, the strength of intraspecific competition (evaluated at the single species equilibrium) and interspecific competition (evaluated at the single species equilibrium of the other species) for both species are equal (Figure 2C and D). Under the scenario where neither species shifts its niche with size (represented by the black dot in Figure 2A) the invasion growth rate of species 1 is highly sensitive to small increases in the competitive ability of larger individuals of species 1 with the majority of the sensitivity being derived from increases in larger individuals (Figure 2E). The sensitivity of the invasion growth rate of species 1 to small increases in niche shifts in species 2 is very small and negative (Figure 2E). For species 2, small increases in the size-dependent competitive asymmetries in species 1 decreased the invasion growth rate of species 2 with the largest effect on the smallest individuals of species 2 (Figure 2F). Small increases in the size-dependent niche shifts of species 2 increased the invasion growth rate of species 2, but this sensitivity was very small across the range of sizes for species 2 (Figure 2F). Together these results suggest that ontogenetic niche shifts alone do not promote coexistence.

**Figure 2.**
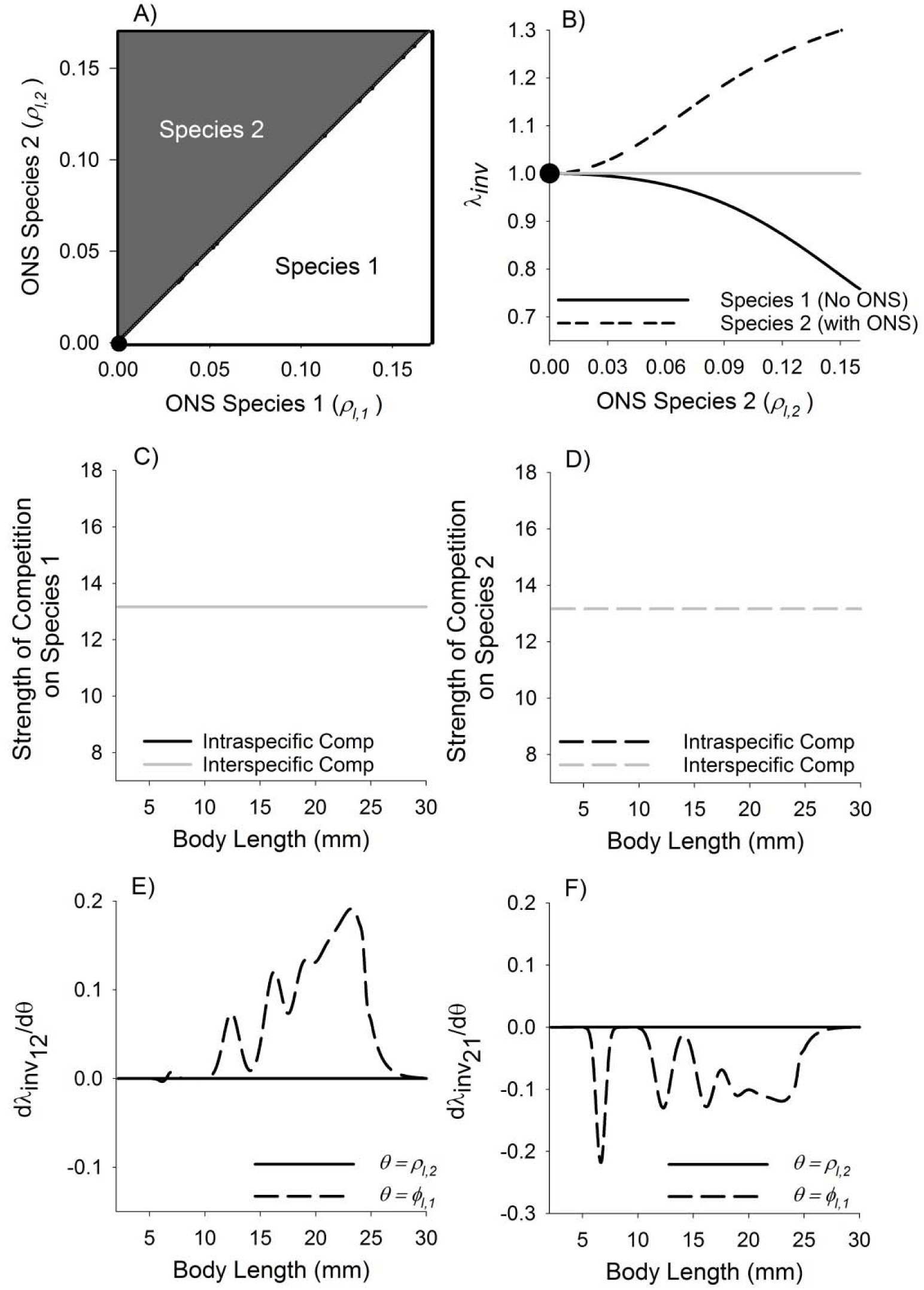
Results of scenario 1 where we examine the coexistence of two species with the same life history and where all individuals of both are equally competitive (competitive symmetry,*ϕ*_*l*,1_ = 0). Panels represent A) coexistence plot, B) invasion growth rate when species 1 has no ontogenetic niche shift and species 2 shift to varying degrees, C and D) the strength of competition at the crossover for species 1 and 2, respectively, and E and F) the sensitivity of the invasion growth rate to changes in niche shifts of species 2 and increased competitive ability of species 1 for species 1 and 2, respectively. In panel A, white regions are combinations of*ρ* values where species 1 is predicted to exclude species 2. Dark grey regions are combinations of values where species 2 is predicted to exclude species 1. In panel B, regions of the functions where *λ_inv_* < 1 indicate where one or both trivial equilibria are stable and mutual invasion is not possible. Sections where the *λ_inv_* > 1 represent regions where the trivial equilibria are unstable and mutual invasion, and therefore coexistence, is possible. For panels C and D, the strength of intraspecific competition is measured at the single species non-trivial equilibrium and the strength of interspecific competition is measured at the trivial equilibrium of the invader when *λ_inv_* > 1 given by the points in panels A and B. For panels E and F, the sensitivities are measured at the trivial equilibrium of the invader when *λ_inv_* > 1 given by the points in panels A and B. The distribution of these sensitivities describes how different sizes contribute to invasion.

### ECOLOGICAL SCENARIO 2: COMPETITIVE ASYMMETRY INCREASES WITH BODY SIZE IN ONE SPECIES

Coexistence between the two species is possible when the competitive ability of one of the species increases with increasing body size and the other species shifts its niche through ontogeny (Figure 3A). When competitive ability of species 1 individuals increases with increasing body size (*ϕ*_*l*,1_ = 0.02), coexistence of the species is predicted when the less competitive species (species 2) shifts its niche to a greater degree than the better competitor (species 1) (Figure 3A). Assuming that species 1 does not shift its niche with body size, coexistence between the species is possible when species 2 shifts its niche to a moderate degree (*ρ*_2_ values between ∼0.0542 and 0.1066; black dots in Figure 3A and B).

**Figure 3.**
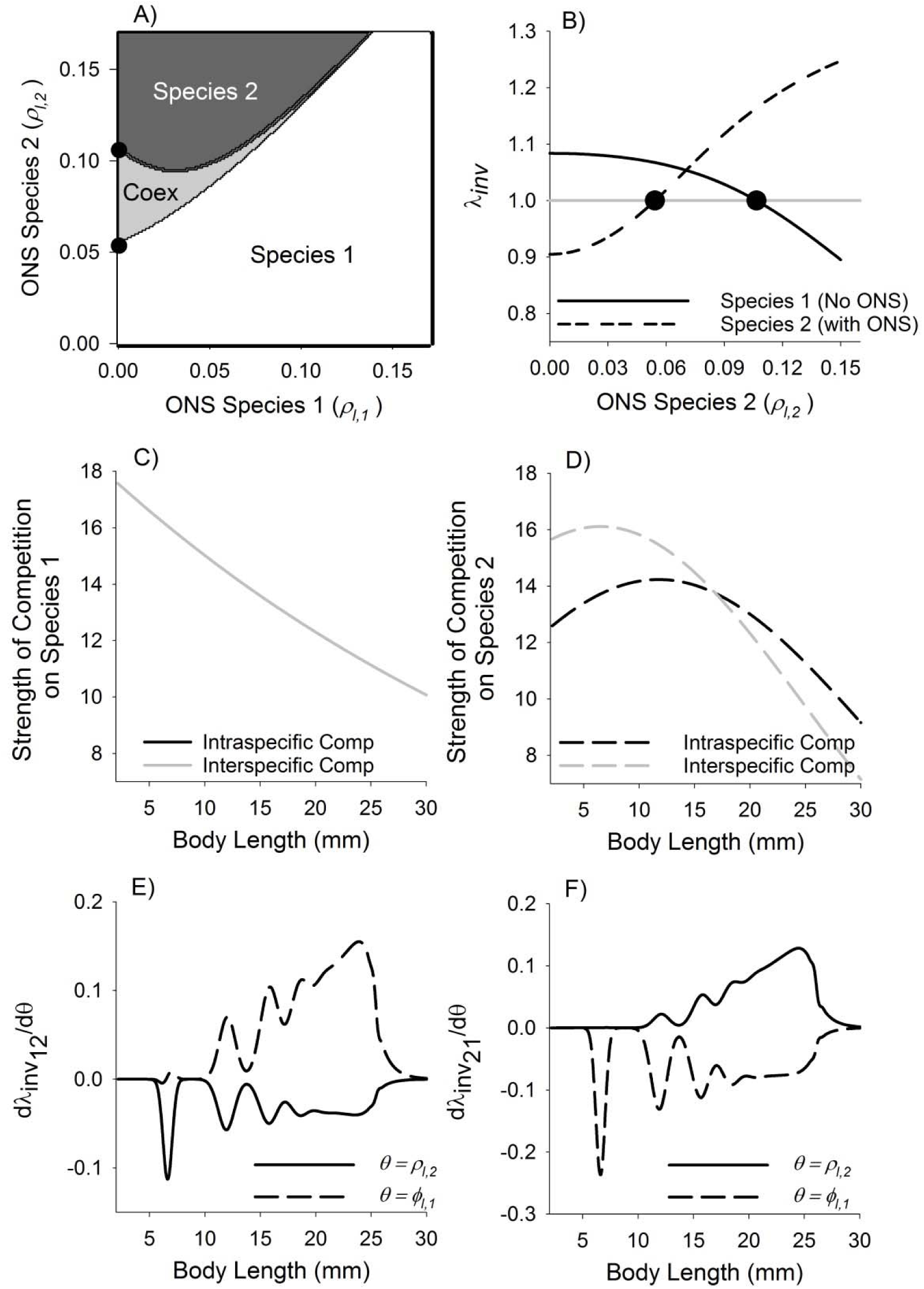
Results of scenario 2 where we examine the coexistence of two species with the same life history and where the larger individuals of species 1 are better competitors (**ϕ_*l*,1_** = **0.02**). Panels represent A) coexistence plot, B) invasion growth rate when species 1 has no ontogenetic niche shift and species 2 shift to varying degrees, C and D) the strength of competition at the crossover for species 1 and 2, respectively, and E and F) the sensitivity of the invasion growth rate to changes in niche shifts of species 2 and increased competitive ability of species 1 for species 1 and 2, respectively. In panel A, white regions are combinations of **ρ** values where species 1 is predicted to exclude species 2. Dark grey regions are combinations of **ρ** values where species 2 is predicted to exclude species 1. Light grey regions are combinations of **ρ** values where the two species are predicted to coexistence (labelled “Coex”). Other panel descriptions are the same as those in Figure 2.

Moving from the origin along the niche shift axis of species 2, coexistence becomes possible when the trivial equilibrium of species 2 becomes unstable (i.e. when *λ*_*inv*21_ becomes greater than 1 in Figure 3B). At this transition point, the strength of intraspecific competition for species 2 (evaluated at the single species equilibrium of species 2) is weaker for smaller individuals and stronger for larger individuals compared to interspecific competition (evaluated at the single species equilibrium of species 1, Figure 3D). Small increases in the niche shift with size in species 2 (*ρ*_2_) is predicted to lead to coexistence (positive derivative in Figure 3F). The effect is concentrated in the largest sizes because these larger sizes are able to escape competition with individuals of species 1. In contrast, small increases in the competitive ability of larger individuals of species 1 (*ϕ*_*l*,1_) decreases *λ*_*inv*_21__ and prevents successful invasion of species 2 (Figure 3F).

Moving further along the niche axis for species 2, a loss of coexistence is then predicted when the trivial equilibrium of species 1 becomes stable (crosses *λ*_*inv*_12__ = 1, Figure 3B). At this point, the strength of intraspecific competition (evaluated at the single species equilibrium of species 1) and interspecific competition for species 1 (evaluated at the single species equilibrium of species 2) decline monotonically with body size and are equal (Figure 3C). For species 1, small increases in niche shift of species 2 prevents invasion by species 1 (Figure 3E). Small increases in the competitive ability of species 1 increases the invasion growth rate mostly through effects on larger individuals of species 1 (Figure 3E). Overall, these results point towards coexistence emerging as a consequence of both ontogenetic changes in resource use and competitive asymmetries.

### ECOLOGICAL SCENARIO 3: COMPETITIVE ASYMMETRY INCREASES WITH SIZE IN SPECIES 1 AND DECREASES WITH SIZE IN SPECIES 2

A wider variety of outcomes is possible when small individuals of species 1 (<15mm) are worse competitors than small individuals of species 2 and large individuals of species 1 (>15mm) are better competitors than large individuals of species 2 (Figure 4). Under this scenario, when species 2 shifts its niche with body size, but species 1 does not (vertical axis in Figure 4A or Figure 5A) the region where coexistence is possible is similar to scenario 2 where coexistence is possible under intermediate levels of ontogenetic niche shifts in species 2 (compare Figure 3A with Figure 4A). However, assuming the each species shifts its niche to an equal and large degree, then each species is able to competitively exclude the other (black areas in Figure 4A or Figure 5A). Which species is able to persist depends on the species that was in the habitat first (priority effect). Below we analyze the model at transition points between these regions and regions of competitive exclusion.

**Figure 4.**
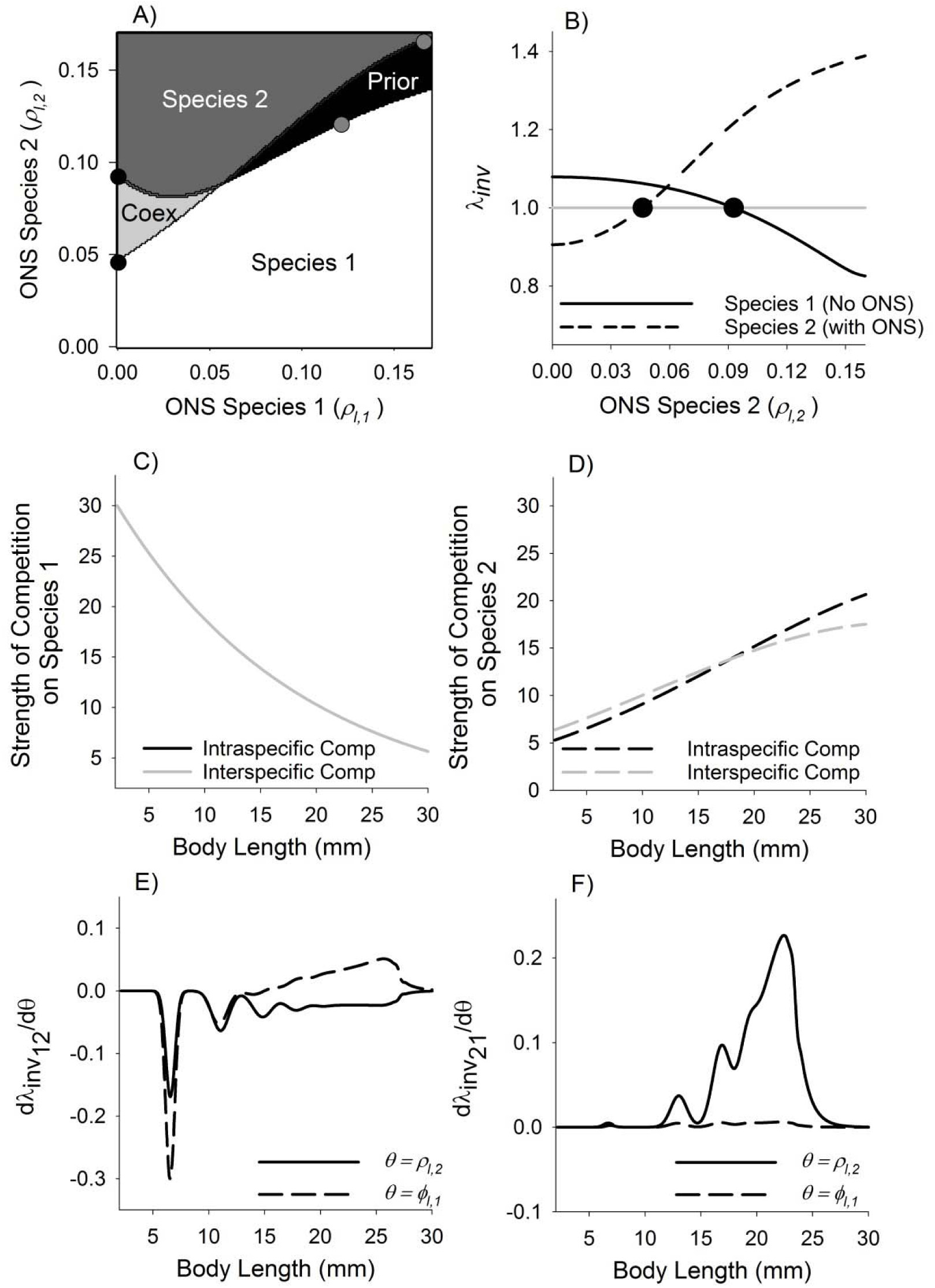
Results of scenario 3 where we examine the coexistence of two species with the same life history and where competitive ability increases with size in species 1 ((**ϕ_*l*,1_** = **0.02**) and decreases with increasing size in species 2 ((**ϕ_*l*,2_** = **−0.02**). Panels represent A) coexistence plot, B) invasion growth rate when species 1 has no ontogenetic niche shift and species 2 shift to varying degrees, C and D) the strength of competition at the crossover for species 1 and 2, respectively, and E and F) the sensitivity of the invasion growth rate to changes in niche shifts of species 2 and increased competitive ability of species 1 for species 1 and 2, respectively. In panel A, white regions are combinations of ***ρ*** values where species 1 is predicted to exclude species 2. Dark grey regions are combinations of ***ρ*** values where species 2 is predicted to exclude species 1. Light grey regions are combinations of ***ρ*** values where the two species are predicted to coexistence (labelled “Coex”). Black regions are combinations of ***ρ*** with priority effects (labelled “Prior”). Other panel descriptions are the same as those in Figure 2.

**Figure 5.**
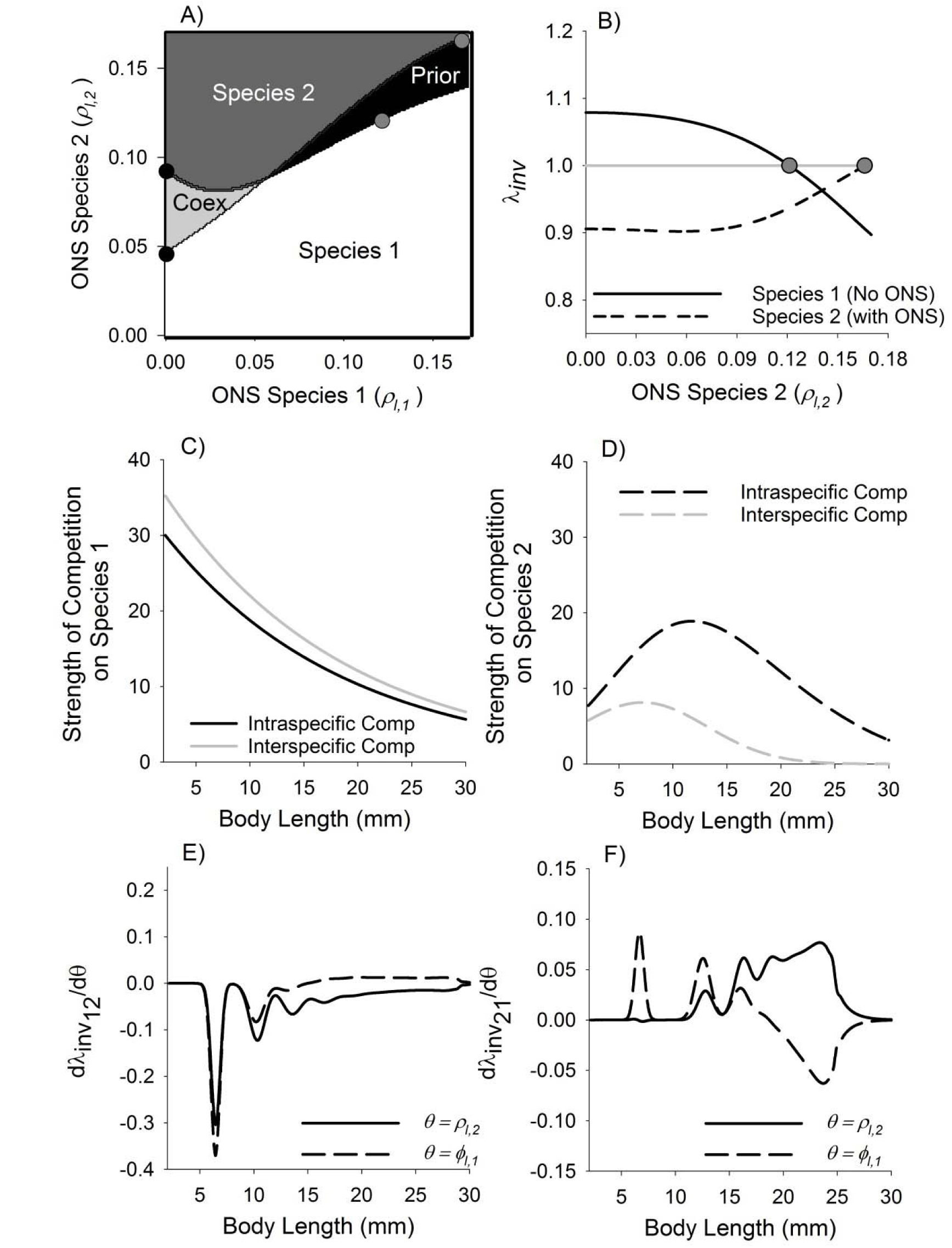
Results of scenario 3 where we examine the coexistence of two species with the same life history and where competitive ability increases with size in species 1 ((**ϕ_*l*,1_** = **0.02**) and decreases with increasing size in species 2 ((**ϕ_*l*,2_** = **−0.02**). Panels represent A) coexistence plot, B) invasion growth rate when species 1 has no ontogenetic niche shift and species 2 shift to varying degrees, C and D) the strength of competition at the crossover for species 1 and 2, respectively, and E and F) the sensitivity of the invasion growth rate to changes in niche shifts of species 2 and increased competitive ability of species 1 for species 1 and 2, respectively. In panel A, white regions are combinations of ***ρ*** values where species 1 is predicted to exclude species 2. Dark grey regions are combinations of ***ρ*** values where species 2 is predicted to exclude species 1. Light grey regions are combinations of ***ρ*** values where the two species are predicted to coexistence (labelled “Coex”). Black regions are combinations of ***ρ*** with priority effects (labelled “Prior”). Other panel descriptions are the same as those in Figure 2.

#### Region of coexistence

Although the region of coexistence in this scenario is similar to that in scenario 2, the underlying reasons for the emergence of this region in this scenario are different. Moving from the origin along the niche shift axis of species 2, coexistence becomes possible when the trivial equilibrium of species 2 becomes unstable (i.e. when *λ*_*inv*_21__ becomes greater than 1 in Figure 4B). At this transition point, and in contrast to scenario 2, the strengths of intraspecific competition (evaluated at the single species equilibrium of species 2) and interspecific competition on species 2 (evaluated at the single species equilibrium of species 1) are roughly equal and increase in strength with increases in body size, indicating a strong influence of density-dependence on larger individuals (Figure 4D). Small increases in the niche shift with size in species 2 (*ρ*_2_) is predicted to lead to coexistence and is almost entirely the result of larger individuals escaping competition with individuals of species 1 (Figure 4F). In contrast, small increases in the competitive ability of larger individuals of species 1 had only a very small positive effect on coexistence (Figure 4F).

Differences in the underlying causes of coexistence also were evident in species 1. At the second boundary along the niche shift axis the trivial equilibrium of species 1 gains stability and species 2 is predicted to competitively exclude species 1 (Figure 4B). At this point, the strengths of intraspecific competition (evaluated at the single species equilibrium of species 1) and interspecific competition on species 1 (evaluated at the single species equilibrium of species 2) are equal and are decreasing functions of size, meaning that competition is strongest in smaller sizes (Figure 4C). Similar to scenario 2, small increases in the ontogenetic niche shift of species 2 decrease the invasion growth rate of species 1 largely through effects on smaller individuals (Figure 4E). Overall, small increases in the competitive ability of large individuals and decreases in the competitive ability of small individuals of species 1 decreases the invasion growth rate of species 1 (sum of derivatives in Figure 4E) with effects through larger individuals increasing and smaller individuals decreasing the invasion growth rate (Figure 4E).

#### Region of priority effects

Moving from the origin along a diagonal where the niche shift in species 1 and 2 are equal (Figure 5A), a region of priority effects emerges when the trivial equilibrium of species 1 becomes stable (*ρ*∼0.1213). At this transition point, the strength of interspecific competition (evaluated at the single species equilibrium of species 1) is stronger than intraspecific competition on species 1 (evaluated at the single species equilibrium of species 2) for all sizes (Figure 5C). Small increases in the ontogenetic niche shifts of species 2 are predicted to decrease the invasion growth rate of species 1 through negative effects on all individuals of species1.

Moving further along the diagonal, the trivial equilibrium of species 2 then becomes unstable at the second boundary along the diagonal (*ρ*∼0.1662). This point approximately represents the situation where small and large individuals of either species do not compete with each other (because they occupy different niches), but where each species is a better competitor in one region of niche space. At this point the strength of intraspecific competition (evaluated at the single species equilibrium of species 2) is much greater than interspecific competition on species 2 (evaluated at the single species equilibrium of species 1, Figure 5D). For intra-and interspecific competition, the strength of competition is hump-shaped across sizes—it is reduced for larger and smaller individuals and is greatest for newborns (interspecific) or individuals of about 12 mm (intraspecific). Small increases in the ontogenetic niche shifts increased the invasion growth rate mostly through effects on larger individuals of species 2 (Figure 5F).

Overall, small increases in competitive asymmetry of species 1 increased the invasion growth rate of species 2 (Figure 5F).

## DISCUSSION

Many theoretical models of coexistence in structured populations envision two discrete life stages (i.e. juveniles and adults) living on two discrete resources (e.g. Haefner and Edson 1984, Loreau and Ebenhoh 1994, Moll et al. 2008, Ackleh and Chiquet 2011). These models are appropriate to model how different types of resource partioning within and between life history stages within and between species facilitate coexistence. However, for many organisms, the resources used, as well as competitive ability, changes throughout ontogeny and it is unclear how predictions from models developed for two stages and two resources overlap with predictions from models explicitly considering continuous change in ontongeny and competitive ability. We developed a discrete time demographic model of two interacting size-structured populations where both the niche and competitive ability can change through ontongeny.

Our results suggest that the routes to coexistence in two interacting species with continuous shifts in the niche are sometimes similar to, and sometimes divergent from, predictions of coexistence of two species where changes in the niche involve discrete changes in life stage. When both species can shift their niche with increasing body size and competition for resources among the sizes is symmetric, then the species that shifts its niche to a greater degree with ontogeny will competitively exclude the other species (Figure 2A). When competitive ability increases with increasing body size, then the two species can coexist when the better competitor shifts its niche with body size to a lesser degree than the weaker competitior (Figure 3A). This region of coexistence shrinks as the better competitor increasingly shifts its niche with increasing size (Figure 3A). When both species shift their niches with size, but each is a better competitor on resources used by smaller or larger individuals, then the model predicts an alternative stable state over some range of niche shifts (Figure 4A and 5A).

Using a two-state, two-resource model in discrete time, Loreau and Ebenhoh (1994) found that when one species shifts its niche with ontogeny and the other does not (vertical axes in our Figures 2-5A) stable coexistence is possible when the shifting species is mainly limited by the stage that escapes competition. When this is not the case, the species that is able to shift its niche outcompete the species that does not. The assumption here is that competition between the species and stages is symmetric. When the species that does not shift is more efficient at utilizing resources (leading to a competitive asymmetry), then the simple life history will competitively exclude the complex life history. These results are different from ours in which the niche can change continuously with increasing body size. Our results show that when one species shifts its niche with body size and the other does not, in the absence of size-dependent competitive asymmetries, the species that shifts its niche will always competively exclude the species that does not shift (Figure 2A). Biologically this is because larger indivdiuals of the shifting species effectively escape competition with individuals that do not shift. We also show that this is the case whenever one species shifts its niche to a greater degree than the other. When competitive ability increases with body size in the species that does not shift its niche is included, then coexistence between the two species is possible if the shifting species does not shift too much. Biological this occurs because bigger individuals in the non-niche shifting species are now imposing more competitive pressure on the smaller individuals of the shifting species. If the niche shifts too much with size, then this effect is weakened.

In the model of Loreau and Ebenhoh (1994), individuals were assumed to completely shift their niche with stage. Schellekens et al. (2010) used a two-state, two-resource bioenergetic model where the niche shift with stage could be complete or incomplete in one species and incomplete in the other. Schellekens et al. assumed resource intake and energetic costs were directly proportional to body size of the consumers. This assumption translates into asymmetries in competitive ability that that are often refered to as relative-size symmetry (Weiner 1990). Thus the overall scenarios modelled are different than the one modelled here where competitive ability is either independent of size or an increasing or decreasing function of size. Our framework is, however, flexible enough to capture the types of changes envisioned in Schellekens et al. (2010).

Adler et al. (2010) constructed a multispecies IPM to examine how intra and interspecies competition in plants influenced coexistence. Their analysis included linear distance between the centers of individual plants as a measure of niche separation. Niche separation was not a function of the size of the individual plants. In our model, we assume that the degree of niche separation can change throughout the life of the individual. This allowed us to capture how competitive interactions can change throughout life. In Adler et al’s model, competitive asymmetries were included by weighting the species-specific competition coefficients by the size of the individual neighbors competing with the focal plant. Such a scaling is similar, but not identical, to the approach here and in Bassar et al. (2016) to model competitive asymmetries. Here and in Bassar et al. (2016) the outcome of competitve interactions were modelled as depending on both the size of the competitor and the size of the focal individual. Scaling to both sizes allows a broader range of competitive interactions to be captured. Moreover, assuming competition scales directly to the size of the competitor only, as in Adler et al., means that competition is relative-size symmetric (Weiner 1990) and essentially that overall density (e.g. number per m^2^) can be replaced with overall biomass density (e.g. grams per m^2^ or whatever metric is used for size). Our flexible framework allows the degree of competitive asymmetry to vary from no asymmetry to situations where larger or smaller individuals are better competitors and allows for a wider variety of density scaling.

The model we present illustrates the variety of outcomes that are possible when both the niche and competitive ability change through ontogeny. We used body size to illustrate the model because it is very often the trait that determines changes in the niche and competitive outcomes. In practice, any underlying continuous trait could be used. Different components of the interaction surface can be eliminated or modified to accommodate a wide variety of niche changes. For example, discrete niche changes based on size could be included by assuming the Gaussian function is one value below a given threshold size and another value above that threshold. Other types of continuous ontogenetic niche changes could be included. For example, instead of the niche shifting its mean position with increases in size, the niche breadth could be modelled as a function of size. This would allow the niches of larger individuals to overlap completely with smaller individuals, but also to occupy some niche regions where they do not compete witih smaller individuals. In other cases, no trait-based competitive asymmetries may exist and so the parameter governing competitive size asymmetries may be safely assumed to be zero.

A key strength of our approach is that our models can be parameterised with data that are routinely collected by ecologists. Fitting the model to data means that researchers can use the model to interpret the results of experiments in a way that integrates all the information on changes in demographic rates as opposed to one demographic rate at a time. Our work, as well as that championed by Adler et al. (2010), illustrate this point, with both research efforts combining a model of demographic rate changes with field data to ask whether competition has a role in structuring biological communities. Our hope is that the framework outlined here will give others the tools to answer questions about coexistence in a wider variety of systems.

The framework presented here can also be used to ask how different competitive interactions between different evolutionary strategies within a species alters natural selection. In this context, aspects of the niche and competitive abilities could be related to body size or any other continuous character. When the trait is body size in the absence of competitive asymmetries, evolutionary strategies that shift their niches with increasing body size would be expected to be favored over those that do not shift their niche. With size based competitive asymmetries, two (or possibly more) strategies may coexist over some range of parameter combinations.

In addition to facilitating tests of evolutionary questions within a single species, the models can be extended to include two or more species each with mulitple evolutionary strategies to ask questions about evolutionary change can promote species coexistence. We parameterized our model using demographic data collected from Trinidadian guppies that live in streams with only one other fish, the killifish (*Rivulus hartii*), with this question ultimately in mind. In Trinidad, guppies also live in streams with more diverse fish communities (Gilliam et al. 1993). These higher diversity streams contain predatory species of fish that significantly decrease the survival rates of guppies compared to streams with only guppies and killifish (Reznick et al. 1996). Decades of research has shown that guppies from these two fish communities display evolutionary changes in life history, behaviour, and morphology (Endler 1980, Reznick and Endler 1982, Reznick 1982, Reznick and Bryga 1987, Reznick 1989, Endler 1995, Reznick et al. 1996, Rodd and Reznick 1997, Gordon et al. 2009) and many of these changes occur within 3-5 years after experimentally introducing guppies into streams that contained only killifish (Reznick et al. 1990). Guppies from these two localities are also know to consume different resources (i.e. they occupy different, but broadly overlapping, resource niches; Bassar et al. 2010, Zandonà et al. 2011, Bassar et al. 2015, El-Sabaawi et al. 2015, Bassar et al. *in press-a*) and may differ in the competitive abilities.

The guppy story is illustrative of what might often occur when a species invades a new habitat devoid of the predators with which it evolved but containing a novel species with which it must compete for resources (Travis et al. 2014). In a deterministic sense, initial invasion and coexistence requires either guppies or killifish to shift their niches with ontogeny and the other to be a better competitor in all sizes or to increase competitive ability with increasing body size. Of course, the precise predictions will depend in part on the structure of the life history of the two species, but so long as these structures are not dramatically different, these predictions are robust. As with guppies and killifish, the invading species and the competitors in the new environment may evolve adaptations that facilitate further coexistence. Both guppies (Reznick 1982, Reznick et al. 1990) and killifish (Gilliam et al. 1993, Walsh and Reznick 2008, 2009, 2010, Walsh et al. 2011, Walsh and Reznick 2011, Furness et al. 2012) are known to evolve when guppies are transplanted into previously guppy-free streams, but how and how much of this evolution is the result of strong interspecific interactions is unknown (Travis et al. 2014, but see Bassar et al. *in press-b*). The signature of these adaptations will be realized in the parameters that describe both the structure of the life history (*α*′*s*) and those that determine the effect of the intra- and interspecific interactions (*ρ*,*ϕ*).

Starting from a set of parameter values that would allow coexistence, evolution could take several routes. Competitive ability could evolve, niche shifts could evolve, body size itself could evolve and therefore the relative importance of different niches would be changing. Predictions regarding the path of least resistence can be identified using the sensitivities of some measure of fitness to the parameters. How the populations respond to these selection pressures involves understanding the variance and covariance structure of the underlying genetic mechanisms. It is overwhelmingly likely that in most cases the underlying genetic architecture of the traits determining these pathways will be quantitative and determined by numerous loci. However, unlike in population genetics where simple demographic models describe the intra and interspecific interactions among different genotypes (MacArthur 1962, Roughgarden 1971, Roughgarden 1976), we know little about how selection on quantitative characters (such as body size, competitive ability, or niche shifts) plays out in an ecological context in species with structured demographic interactions (Childs et al. 2016, Coulson et al. *in review*) and resultingly have little ability to make predictions about the outcome of such coevolutionary interactions. Integrating ecological models such as the ones we describe here with models incorporating genetic mechanisms will undoubtedly yield valuable insights into understanding how evolution helps to shape coexistence and biodiversity, just as it has in the past in evolutionary population genetics. The models and results that we present here represent one step in this direction and our future efforts will be aimed at linking these models with those that describe the genetic mechanisms of quantitative characters and explicitly link together the causes and consequences of ecological and evolutionary processes.

## ACKNOWLEDGEMENTS

We wish to thank Yuridia Reynoso and the numerous field and laboratory technicians that helped with the mesocosm experiments and dissection of the guppies. We also wish to thank Mark Rees, Dylan Childs and Shripad Tuljapurkar for discussions regarding competitive asymmetries. The mesocosm and field research was originally funded by USA NSF research grants (EF0623632, 9419823) and the model development and analyses presented here were funded by a UK NERC grant (ATR00350) and a USA NSF grant (DEB 1556884). TC also acknowledges support from an ERC advanced grant (LEED, number 249872).

## SUPPLEMENTAL MATERIAL FOR

### MATRIX APPROXIMATION OF THE IPM

As is typical for IPM’s, the integro-difference equation describing the per time step dynamics (equation 1) is converted to a matrix approximation prior to analysis (Easterling et al. 2000). The matrix approximation of the IPM for species *i* is then:

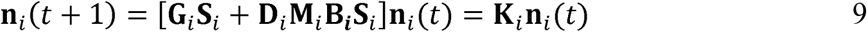
where **n**_*i*_(*t*)is a vector containing the density of individuals in the population of species *i* with trait value *z* at time *t*. Each vital rate matrix still depends on the density of each species at time *t*, but for simplicity we have dropped the parentheses. We used matrices of dimension 200 x 200 in all analyses and used the midpoint of each size class in our approximation.

### STABILITY OF THE TRIVIAL EQUILIBRIUM

The equation describing the dynamics of species *i* at the trivial equilibrium of species *i* and the single species equilibrium of species *j* is:

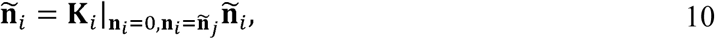
Where 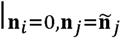 means that the matrix is evaluated at the population distribution of species *i* equal to zero for all sizes and at the single species equilibrium of species *j*. The stability analysis proceeds by applying a small perturbation to the population size vector of species *i* such that:

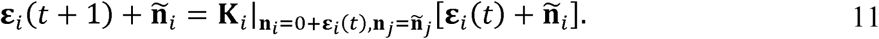
Converting to vector format and expanding 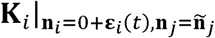 in a first order Taylor series around **n**_*i*_ = 0 results in:

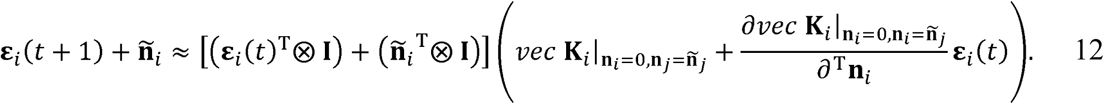

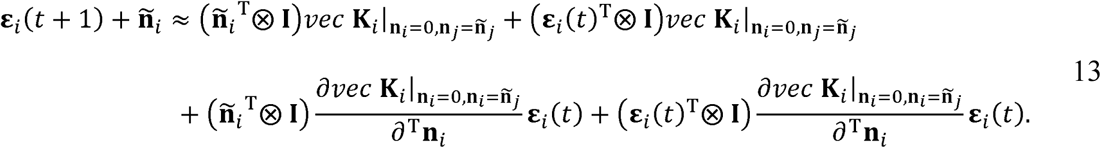

The first term is equal to **ñ**_*i*_ and cancels with **ñ**_*i*_ on the left side of the equation. The third term is equal to zero because **ñ**_*i*_ = 0. The last term is of second order and is ignored when entries in **ε**_*i*_(*t*) are small. Making these changes leaves:

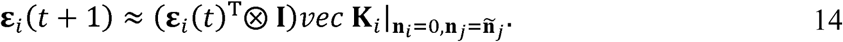

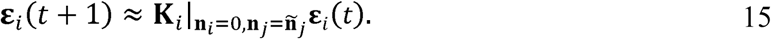

The stability of the trivial equilibrium is then given by the dominant eigenvalue of the matrix 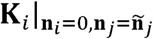 (Caswell 2001). This eigenvalue is also the invasion growth rate, *λ_inv_ij__*, or the eventual growth rate of the invading species. Very importantly, in this context 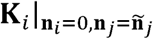 measures deviations, **ε**_*i*_, not individuals. This means that the equation is linear even though equation 10 is non-linear.

### SENSITIVITY ANALYSIS OF THE TRIVIAL EQUILIBRIUM OR INVASION GROWTH RATE

If **p**_*i*_is a right eigenvector corresponding to*λ_inv_ij__* and **v**_i_^T^ is the corresponding left eigenvector, where **v**_*i*_^T^**K**_*i*_ = *λ_inv_ij__***v**_*i*_^T^ and **v**_*i*_^T^**P**_*i*_ = 1, then

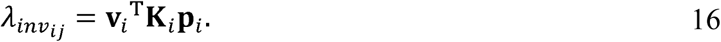

Note that for brevity, we have dropped the 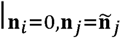 from 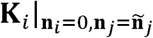. The derivative of *λ_inv_ij__* with respect to **K**_*i*_ is:

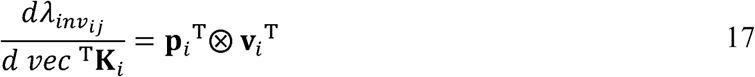
where ⊗ means to take the kronecker product. Because equation 15 is linear, we need not additionally consider the derivatives describing how **p**_*i*_ and **v**_*i*_ change with changes in **K**_*i*_. In a typical density-dependent (non-linear) model, these derivatives ensure that the sum of the total derivative, 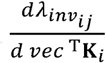, is always equal to zero. To get how any underlying parameter changes *λ_inv_ij__* multiply equation 17 through by 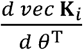 to get:

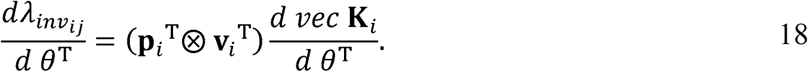

Recognizing that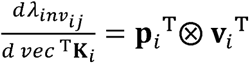, rewrite as:

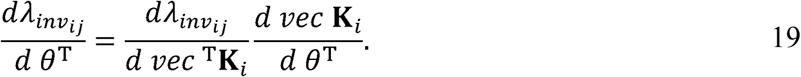

Now any parameter can change **K***_i_* directly because **K***_i_* is a function of *θ* and indirectly through changes in **n**_*i*_ or **n**_*j*_. Expanding 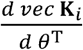 to capture this yields:

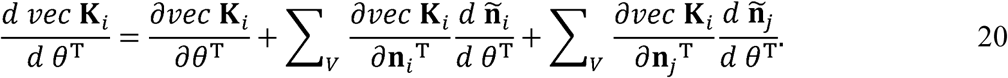

The first term is the direct effect of the parameter on **K***_i_*. The second and third terms are the indirect effects of the parameter(s) on **K***_i_* through how they change the population size distribution of species *i* and species *j*, respectively. The summations mean to sum the derivatives across all the vital rates. This is done because the although the parameter may not be a parameter of a given vital rate, it can still alter the vital rate indirectly through its effect on the population distribution. Inserting this into equation 19:

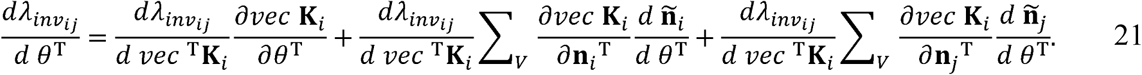

The next step involves calculating the effect of the parameters on the equilibrium population sizes of both species. These are:

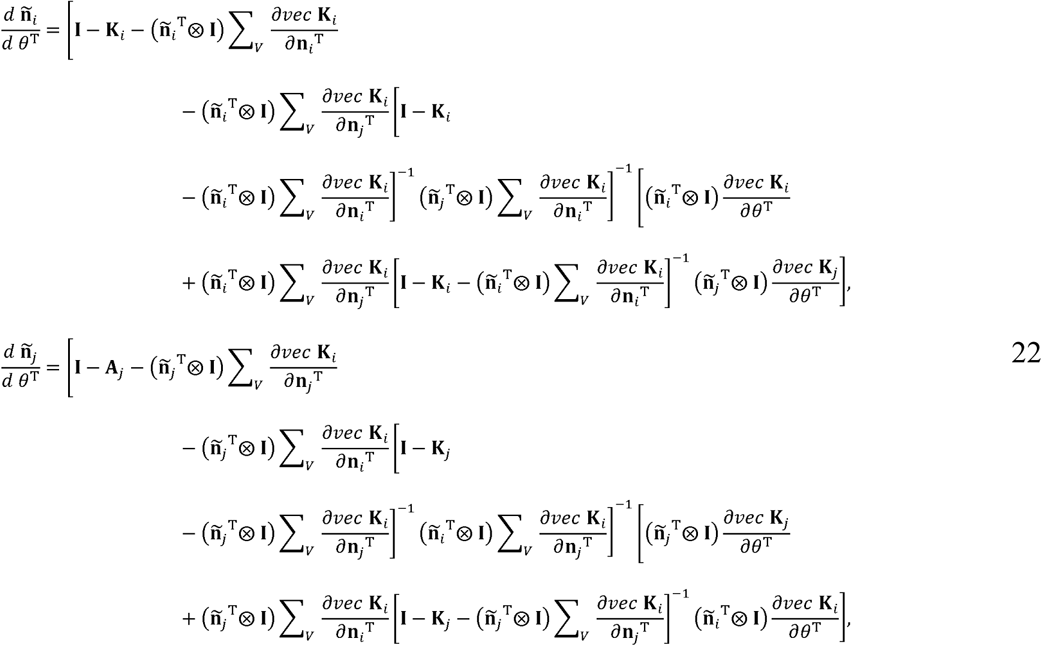

Inserting these into equation 21 makes it in terms only of the parameters and their effects through changing the population size distributions. These are, of course, horrid equations, but some terms can be eliminated by assumption. These terms are any term with **ñ**_*i*_ because we assumed. *N_i_* = 0. Therefore equation 22 becomes:

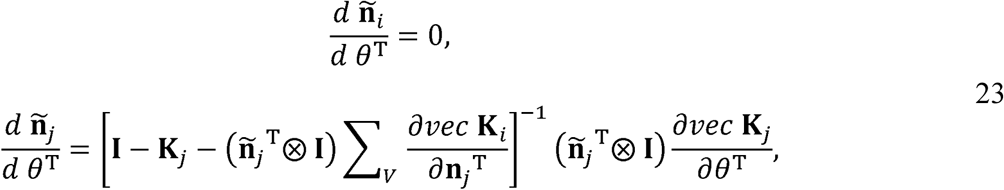
and so equation 21 becomes:

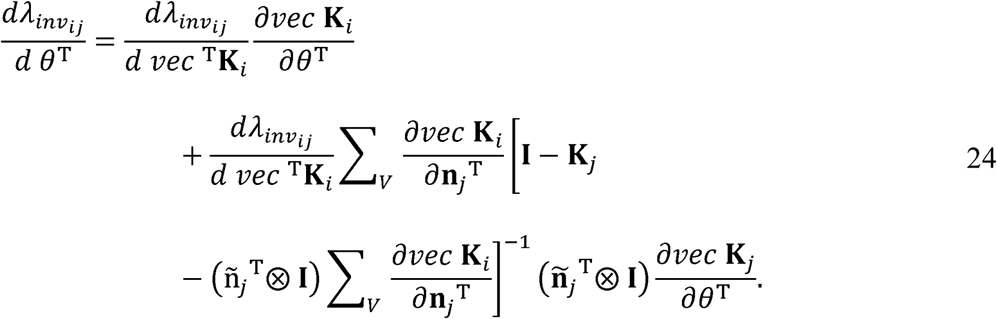

This is a full accounting of how changing a parameter(s) can influence invasion success through any or all of the demographic rates.

When *θ* is a parameter defining the interaction surface, we can expand the derivatives 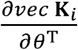 and 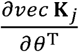 using the chain rule to reveal the underlying pathways. These expansions are:

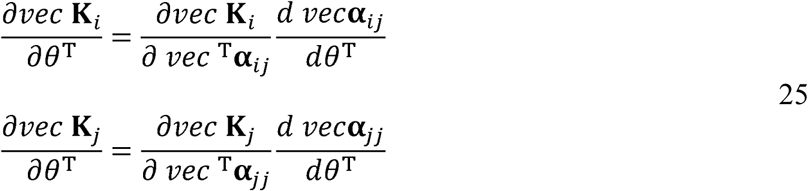

The expansion of 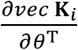 yields one term describing how the projection matrix changes through interspecific interactions with species *j*. The intraspecific interaction term within species *i* ultimately drops out because using the invasion framework *N_i_* = 0. The expansion of 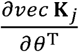 yields one term that describes how changes in the parameter changes intraspecific interactions in species *j*. The interspecific term for species *j* is not presented because it will also drop out because N_*i*_ = 0. Substituting these expansions into equation 24 and doing a little rearranging:

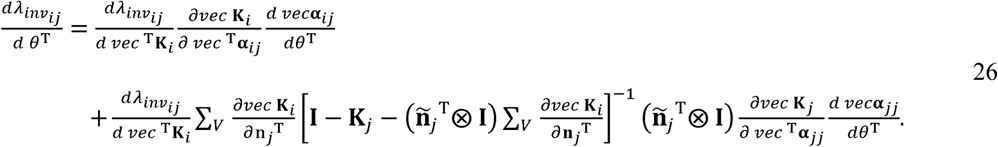

Making 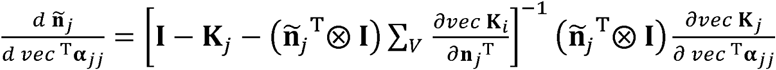 simplifies the notation a bit so that:

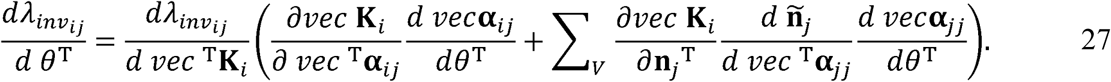

Now the equation is separated into terms that describe how the parameter changes intra and interspecific competition. The first term describes how interspecific competition influences invasion directly through changes in the parameter. The second terms describes how changes in the parameter alter the equilibrium population distribution of species *j*.

Because equation 27 represents the inner product of two vectors, it results in a scalar that incorporates all the effects a change in *θ* across all sizes. Our goal is to keep the effects on *λ_inv_ij__* through different sizes separate. To do so, rewrite equation 27 so that:

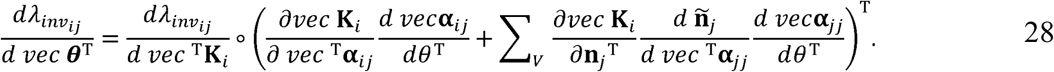
where ◦ means to take the Hadamard product. Equation 28 is a vector with the form:

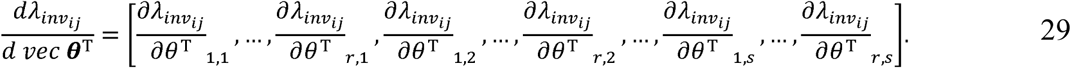

It is the vectorization of a matrix of partial derivatives with dimensions that correspond to the dimensions of **K***_i_* and row elements *r* and column elements *s*. Summing all elements of equation 28 is equal to equation 27. Summing all *r* row elements for any *s* column, thus yields the sensitivity of *λ_inv_ij__* to *θ* through size *s* individuals:

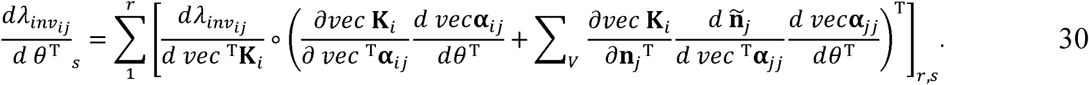

